# The Scar/WAVE complex drives normal actin protrusions without the Arp2/3 complex, but proline-rich domains are required

**DOI:** 10.1101/2022.05.14.491902

**Authors:** Simona Buracco, Shashi Singh, Sophie Claydon, Peggy Paschke, Luke Tweedy, Jamie Whitelaw, Lynn McGarry, Peter A. Thomason, Robert H. Insall

## Abstract

Cell migration requires the constant modification of cellular shape by reorganization of the actin cytoskeleton. The pentameric Scar/WAVE regulatory complex (WRC) is the main catalyst of pseudopod and lamellipodium formation. Its actin nucleation activity has been attributed to its ability to combine monomeric actin and Arp2/3 complex through the VCA domain of Scar/WAVE, while other regions of the complex are typically thought to mediate spatial and temporal regulation and have no direct role in actin polymerization.

Here we show that the Scar/WAVE with its VCA domain deleted can still induce the formation of morphologically normal actin protrusions. Equivalent results are seen in B16-F1 mouse melanoma cells and *Dictyostelium discoideum* cells. This actin polymerization occurs independently of the Arp2/3 complex, whose recruitment to the leading edge is greatly reduced by the loss of the VCA domain. We also expressed Scar/WAVE with VCA and polyproline domains both deleted. In *Dictyostelium* cells, these were only active if WASP (which contains its own proline-rich domain) was available. Similarly, in B16-F1 cells both Abi and WAVE proline-rich domains needed to be deleted before the function of the WRC was lost. Thus we conclude that proline-rich domains play a central role in actin nucleation.

Our data demonstrate a new actin nucleation mechanism of the WRC that is independent of its VCA domain and the Arp2/3 complex. We also show that proline-rich domains are more fundamental than has been thought. Together, these findings suggest a new mechanism for WRC action.

## Introduction

Actin polymerization is an essential mechanism for multiple processes in eukaryotic cells, including cell migration, cytokinesis and vesicle trafficking^1-4^. Its many regulators include the members of the Wiskott-Aldrich syndrome protein (WASP) family. These proteins are highly conserved throughout evolution and share a similar domain structure: an N-terminal WH1 domain, a central proline rich region (here referred to as polyproline domain or PP domain) and a C-terminal VCA region^5^. Thanks to its ability to bind both actin monomers (via the V region, also known as WASP homology 2 or WH2) and the Arp2/3 complex (via the CA region), the VCA domain can promote actin nucleation by the Arp2/3 complex and induce the formation of branched actin networks^6,7^. Indeed, the actin nucleation activity of WASP-family proteins is traditionally attributed exclusively to their C-terminal region, while the N-terminal parts are expected to mediate spatial-temporal regulation and have no direct role in actin polymerization.

Among the WASP-family proteins, WAVE (WASP family Verprolin homolog — also known as SCAR for suppressor of cAMP receptor) regulation is exceptionally complex due to the protein being constitutively incorporated into a large hetero-pentamer of ∼400 kDa, known as the WAVE regulatory complex (WRC). The five subunits (namely Nap1/NCKAP1, PIR121/CYFIP, Scar/WAVE, Abi and HSPC300/Brk1) interact forming an autoinhibited structure that sequesters WAVE VCA domain within the complex. According to most current models, when a signal triggers its activation, the WRC undergoes a conformational change that exposes the VCA domain to activate Arp2/3^8^. This induces the formation of the branched actin networks that form cell protrusions and promote cell migration. Cells lacking a functional WRC complex struggle to make protrusions and their movement is profoundly compromised^9-11^.

In the last decade there have been a few studies that suggested that the VCA domain is not the only section of WASP family proteins involved in actin polymerization. In vitro studies proved that the addition of the polyproline region to the VCA domain enhances the rate of filament elongation both in the presence and in the absence of the Arp2/3 complex^12,13^. In the budding yeast *Saccharomyces cerevisiae* the polyproline domain of the WASP orthologue Las17 nucleates actin filaments in the absence of the VCA domain and independently of Arp2/3^14,15^. Moreover, a recent paper demonstrated that Arp2/3-null cells still form lamellipodia (sheet-like actin protrusions) where the WRC complex localizes to the protruding edge^16^. Emerging from these observations is the idea that WASP-family proteins could support actin nucleation through effectors other than the Arp2/3 complex, and their PP domains may play a central role in this newly-discovered activity. However, so far no similar functions have been associated to the PP domain of WASP-family proteins in any other organism aside from yeasts.

Here, in order to achieve a better understanding of the different contributions of PP and VCA regions to actin nucleation, we expressed truncated forms of the Scar/WAVE protein with either the VCA domain, or both VCA and PP domains deleted in two different cellular models known for their high motility: B16-F1 melanoma cells and the amoeba *Dictyostelium discoideum*. Our data demonstrate that the VCA domain is dispensable for cell migration, and a WRC complex lacking its VCA region is capable of catalyzing actin polymerization in cell protrusions. We also prove that PP domains are key regulators of pseudopod and lamellipodia formation, and suggest an actin-polymerization function of the WRC complex that is independent of Arp2/3.

## Results

### WAVE2 VCA domain is not essential for lamellipodia formation

We generated a construct for the expression of WAVE2 lacking its VCA domain (WAVE2ΔVCA, Fig.1A) and expressed it in B16-F1 mouse melanoma cells. These cells are normally highly motile and able to form broad stable lamellipodia. To avoid the compensatory effect of the native proteins, we used an engineered B16-F1 cell line in which the two genes encoding WAVE1 and WAVE2 have been disrupted (WAVE1/2 KO). As a result, these cells are unable to generate lamellipodia and poorly migrate using filopodia-like protrusions^11^. These defects can be easily rescued by the expression of WAVE2 alone^17^ (here indicated as WAVE2 full length or WAVE2 FL, Fig.1A). By western blot we confirmed that expression of WAVE2ΔVCA did not affect the levels of other WRC members, as compared to WAVE2 FL expressing cells (Fig.1B). Moreover, pulldown of Cyfip1 and NCKAP1 using an EGFP-tagged version of WAVE2ΔVCA as bait confirmed its inclusion in the complex (Fig.1C).

**Fig. 1.**
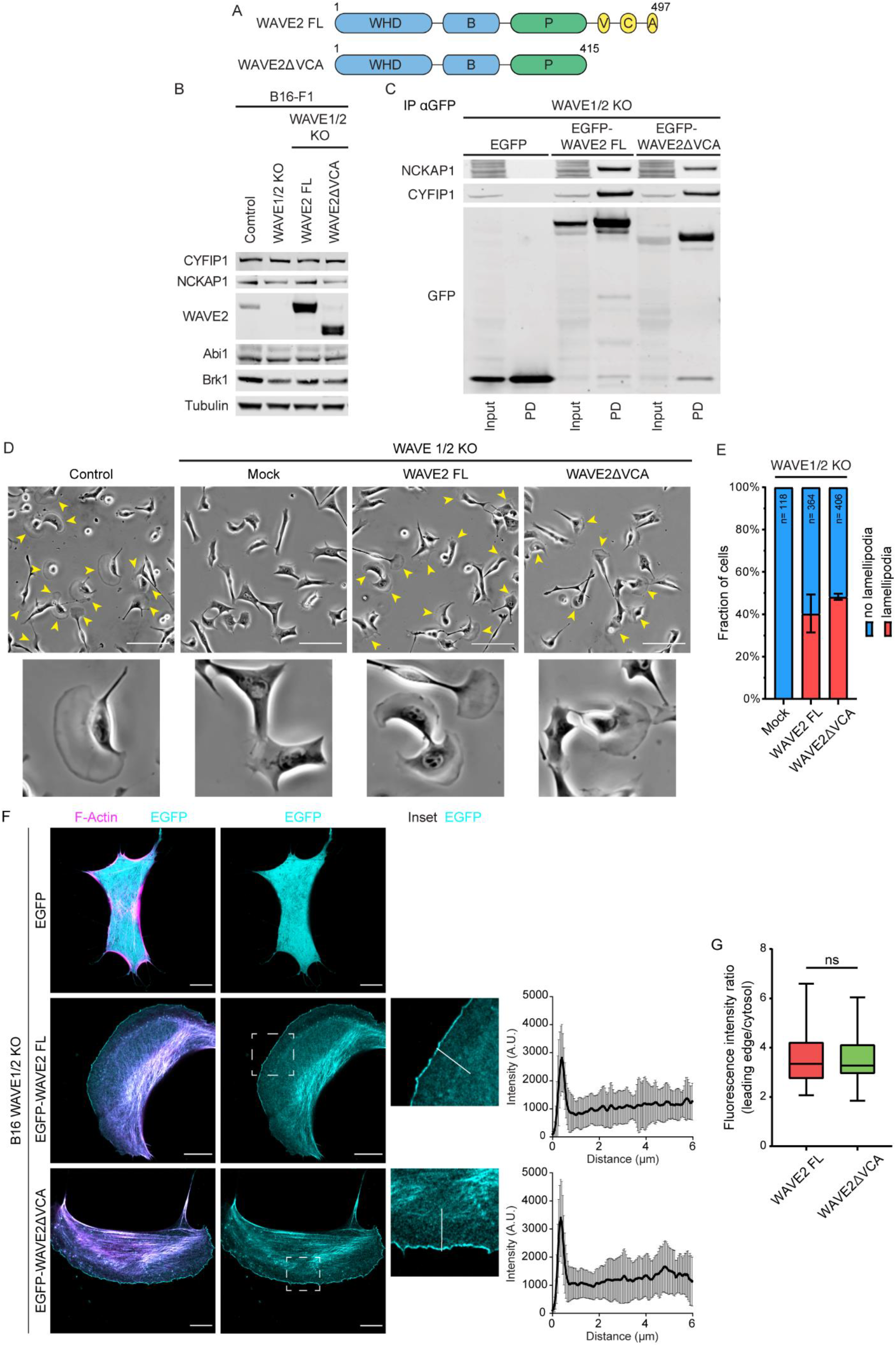
WAVE2ΔVCA rescues the formation of lamellipodia in WAVE1/2 KO cells. **(A)** Schematic of mouse WAVE2 FL and WAVE2ΔVCA showing amino acid numbers and domains. WHD – WASP homology domain; B – basic domain; P – polyproline domain; V – verprolin homology region; C – central region; A – acidic region. **(B)** Representative western blot of lysates of B16-F1 cells, WAVE1/2 KO cells, as well as KO cells expressing WAVE2 FL or WAVE2ΔVCA to detect expression levels of WAVE complex components, as indicated. Tubulin was used as loading control. **(C)** Western blot of GFP immunoprecipitation from lysates of WAVE1/2 KO cells expressing either EGFP alone, EGFP-WAVE2 FL or EGFP-WAVE2ΔVCA, probed with anti-NCKAP1 and anti-Cyfip1. **(D)** Rescue of lamellipodia formation in B16-F1 or WAVE1/2 KO cells transfected with WAVE2 FL or WAVE2ΔVCA and plated on laminin-coated 6-well plates. Yellow arrows indicate lamellipodia protrusions and insets below show magnifications of single cells. Scale bar = 100 µm. **(E)** Quantification of cells in **D** presenting with or without lamellipodia as a percentage. Error bars represent SD, *n* > 118 cells counted for each condition from 2 independent experiments. **(F)** WAVE1/2 KO cells were transfected with LifeAct-TagRed (magenta) and EGFP-tagged WAVE2 (cyan) constructs as indicated, and plated on laminin-coated coverslips for analysis of lamellipodia morphology and localization of the WRC. Representative cells are shown. The inset depicts a zoomed in area used for the quantification of EGFP-WAVE2 intensity. Graphs on the side show the quantification of fluorescence intensity along the white line with 0 corresponding to the leading edge of the lamellipodia. The scale bar represents 10 µm. Error bars represent S.D. *n*> 21 cells counted for each condition from 3 independent experiments. **(G)** Quantification of WAVE2 recruitment as ratio between fluorescence intensity at the leading edge and in the cytosol. Bars show min to max values. Statistical significance was assessed by a two-tailed *t*-test. NS, not significant.

Next, we explored the ability of WAVE2ΔVCA transfected B16-F1 cells to form lamellipodia. According to current models, a WRC complex lacking its VCA domain should be unable to interact with the Arp2/3 complex, and thus unable to promote actin nucleation. However, WAVE2ΔVCA expression restored the formation of lamellipodia in WAVE1/2 KO cells migrating on laminin (Fig.1D). The rate of lamellipodia rescue was comparable to that induced by the expression of WAVE2 FL, with >40% of cells able to form lamellipodia when expressing either WAVE2 FL or WAVE2ΔVCA (Fig.1E). This result suggests that the WRC lacking its VCA domain can still promote the nucleation of the branched actin network that is the principal constituent of cellular lamellipodia. One limitation to our understanding of WRC function is that there are no direct assays to identify the activated WRC and distinguish it from the autoinhibited cytoplasmic form. Currently, lamellipodial formation and recruitment of the WRC to the membrane at the leading edge of protrusions is the best indicator of its activation. According to this indicator, a direct role for WAVE2ΔVCA in lamellipodial generation was further supported by the protein localization inside the cell: like the full-length protein, EGFP-WAVE2ΔVCA was recruited to the leading edge of protrusions, where it accumulated in a sharp and continuous line (Fig.1F). Quantification of its fluorescence intensity confirmed that the truncated WRC is recruited to the plasma membrane at comparable levels to the control containing WAVE2 FL (Fig.1G).

### WAVEΔVCA promotes actin polymerization through an Arp2/3-independent mechanism

Since no differences were observed in the ratio of lamellipodia formation promoted by WAVE2 FL and WAVE2ΔVCA, we looked for possible morphological alterations caused by the deletion of the VCA domain. B16-F1 cells expressing EGFP as control, or either EGFP-WAVE2 FL or EGFP-WAVE2ΔVCA, were seeded on laminin-coated plates and stained with phalloidin. A systematic analysis of their morphological properties was carried out using high content imaging. Based on the pattern of F-actin and on cell shape, we separated the transfected cells into three classes: no lamellipodia, forming a mature lamellipodia or forming immature lamellipodia (identified as small, narrow, and tipped by filopodia) (Fig.2A). About 19% of cells expressing WAVE2 FL were classified as forming mature lamellipodia. This percentage lowered to 10% for cells expressing WAVE2ΔVCA. The proportion of cells forming immature lamellipodia was comparable between the two conditions. However, when considering only those cells with lamellipodia, the analysis reveals a higher percentage of immature lamellipodia in WAVE2ΔVCA expressing cells than is seen in those cells expressing the full-length protein (Fig.2B). This difference was further confirmed by the quantification of lamellipodia width: cells rescued by WAVE2ΔVCA made smaller protrusions compared to WAVE2 FL expressing cells (Fig.2C). Thus, a WRC deprived of its VCA domain is still able to promote the formation of lamellipodia, but a higher percentage of them are small and immature, suggesting a reduction of robustness.

The increased rate of immature protrusions could indicate a defect in actin polymerization. Since the Arp2/3 complex is the major actin nucleator of lamellipodia, we quantified its recruitment at the plasma membrane of protrusions. As expected, Arp2/3 complex immunostaining is clearly visible at the leading edge of WAVE2 FL expressing cells, consistent with their ability to form fully developed lamellipodia. However, Arp2/3 recruitment at the plasma membrane was strongly reduced in cells expressing WAVE2ΔVCA (Fig. 2D-E). Moreover, only WAVE2 FL and not WAVE2ΔVCA could co-precipitate the Arp2/3 complex (Fig. 2F). Without the VCA domain the WRC is thus unable to bind to the Arp2/3 complex, and this results in a reduction of Arp2/3 recruitment to the leading edge.

**Fig. 2.**
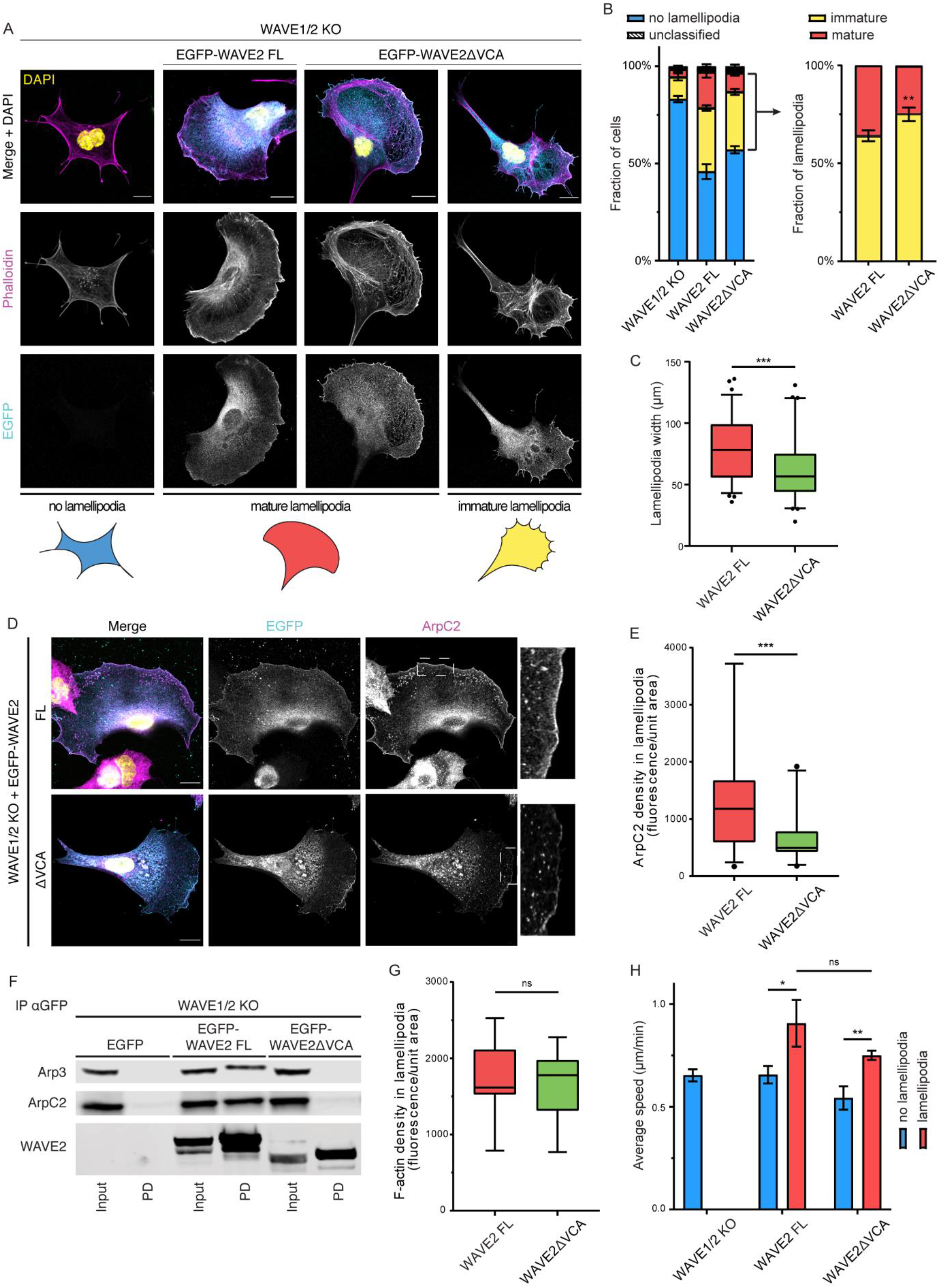
Morphological analysis of lamellipodia generated by WAVE2ΔVCA-expressing cells. **(A)** Cell morphologies and lamellipodia phenotypes of WAVE1/2 KO cells transfected with EGFP, EGFP-WAVE2 FL or EGFP-WAVE2ΔVCA, and stained for the actin cytoskeleton with phalloidin. Cells were classified as not forming lamellipodia (in blue), forming immature lamellipodia (in yellow) or forming mature lamellipodia (in red). Representative cells for each phenotype are shown. Scale bars = 10 µm. **(B)** Quantification as a percentage of morphological classification of cells as in **A** using an High-Content Screening system. Inset represents only cells that present with lamellipodia, grouped as mature or immature as a percentage. Error bars represent S.D., *n* > 1500 cells counted for each condition. **(C)** Quantification of lamellipodial width measurements. Bars show 5-95 percentile, *n* > 63 cells counted for each condition. **(D)** Representative WAVE1/2 KO cells expressing EGFP, EGFP-WAVE2 FL or EGFP-WAVE2ΔVCA and stained for the Arp2/3 complex subunit ArpC2. EGFP is shown in cyan. The protein of interest is shown in magenta and an inset on the side shows the leading edge localization of the protein of interest. Scale bars = 10 µm. **(E)** Quantification of ArpC2 intensity at lamellipodia. Bars show 5-95 percentile, *n* > 21 for each condition. **(F)** Western blot of GFP immunoprecipitation from lysates of WAVE1/2 KO cells expressing either EGFP alone, EGFP-WAVE2 FL or EGFP-WAVE2ΔVCA, probed with anti-Arp3 and anti-ArpC2 for identification of corresponding subunits of the Arp2/3 complex. **(G)** Quantification of F-actin intensity levels in the lamellipodium obtained from phalloidin stainings. Bars show min to max values, *n* > 13 for each condition. **(H)** Random migration assay with WAVE1/2 KO cells expressing WAVE2 FL or WAVE2ΔVCA, and analyzed as described in methods. Cells with and without lamellipodia are displayed separately. Graph shows mean values from 3 independent experiments. Error bars represent S.D. For quantifications in **C, E, G** and **H** statistical significance was assessed by a two-tailed *t*-test. NS, not significant, * p < 0.05, ** p < 0.001, *** p < 0.0001.

Reduced Arp2/3 complex is expected to correspond to a reduction of actin polymerization in protrusions. To investigate if this holds true in our system, we quantified lamellipodial F-actin intensity in phalloidin-stained cells. Surprisingly, no differences were identified between WAVE2 WT and WAVE2ΔVCA expressing cells, suggesting no alteration in the rate of actin polymerization (Fig.2G). These results translated into similar efficiency of random migration: both truncated and full length WAVE2 rescued the migration speed of WAVE1/2 KO cells (the analysis was split between cells with and without lamellipodia because of the large differences in migration efficiency between these two conditions, Fig.2H). Therefore, WAVE2ΔVCA domain can still promote actin polymerization and rescue B16-F1 motility. This result together with the reduction of Arp2/3 recruitment suggests that WAVE2ΔVCA actin polymerization activity is uncoupled from the Arp2/3 complex. However, the increased rate of immature lamellipodia may indicate that this alternative mechanism differs in its regulation of actin dynamics, and may favour less stable protrusions.

### *Dictyostelium* ScarΔVCA can rescue pseudopod formation and cell motility

To test whether the VCA-independent actin nucleating activity of the WRC is conserved throughout evolution, we used an equivalent approach in *Dictyostelium discoideum* cells. This model organism is ideal to study the WRC. The primary structures of its subunits are well conserved between species (mammalian CYFIP1, NCKAP1, Brk1 and Abi are all highly similar to their *Dictyostelium* paralogues^18^). Moreover, the complex members are all encoded by single genes, and the haploid genome makes it extremely easy to genetically target them^18-21^. We generated a plasmid for the expression of Scar and removed its VCA domain (ScarΔVCA, Scar is here used in place of WAVE to distinguish between mammalian and *Dictyostelium* homologs, Fig3A). Applying the same approach previously described for B16-F1 cells, we expressed ScarΔVCA in a *Dictyostelium* Scar KO cell line (Fig.3B). As a control, we used a plasmid for the expression of unmutated full length Scar (Scar FL), whose expression rescued the migration defects of Scar KO cells^17^. Like Scar FL, ScarΔVCA was proved to be correctly incorporated into the WRC complex by immunoprecipitation (Fig.3C). However, deletion of Scar VCA domain destabilized the complex, with ∼70-80% reduction of the levels of all other members of the WRC compared to Scar FL expressing cells (Fig.3B, Supplementary Fig.1A-D).

We then tested the ability of ScarΔVCA to rescue parental phenotype in Scar KO cells. Cells lacking Scar migrate mainly by blebbing, and can generate relatively rare pseudopods, which results in an inefficient and less directional migration^22^. Similar to Scar FL, expression of ScarΔVCA was able to rescue the ability of cells to form pseudopods (Fig.3D). Like in B16-F1 cells, in *Dictyostelium* active WRC complex localizes at the leading edge of migrating cells, where it causes the formation of actin protrusions^22^. Using GFP-Nap1 as a reporter for the complete WRC, we observed that both Scar FL and ScarΔVCA rescued the localization of the complex at the pseudopod periphery, where it accumulated in a continuous line at the leading edge followed by a patch of enriched actin (Fig.3D). Surprisingly, despite the higher instability of the truncated complex, no differences in the recruitment levels to the protruding edge were identified between Scar FL and ScarΔVCA containing WRC (Fig 3E). The motility defects of Scar KO cells were also partially rescued by ScarΔVCA expression, with recovery of about 50% of both speed, chemotaxis efficiency index and directedness (Fig.3F-H). Thus, the ability of Scar/WAVE to promote the formation of actin protrusions even after deletion of its VCA domain is not limited to a mammalian system, like B16-F1 cells, but is conserved in *Dictyostelium* cells.

**Fig. 3.**
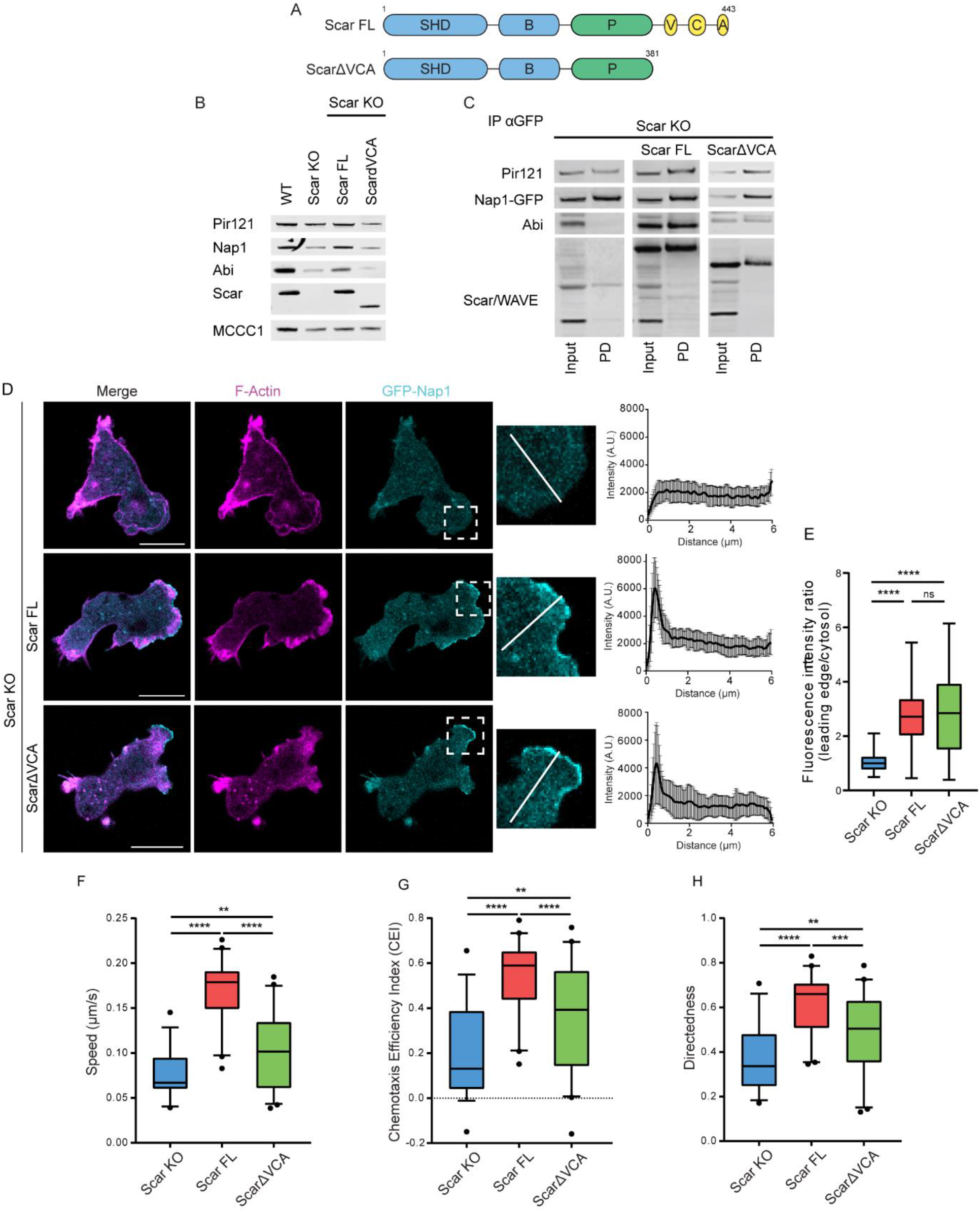
Expression of ScarΔVCA rescues motility and pseudopod formation in *Dictyostelium* Scar knockouts. **(A)** Schematic of *Dictyostelium* Scar FL and ScarΔVCA showing amino acid numbers and domains. SHD – Scar homology domain; B – basic domain; P – polyproline domain; V – verprolin homology region; C – central region; A – acidic region. **(B)** Representative western blot of lysates of *Dictyostelium* AX3 WT cells, Scar KO cells, as well as KO cells expressing Scar FL or ScarΔVCA to detect expression levels of WRC components, as indicated. MCCC1 was used as loading control. **(C)** Western blot of GFP immunoprecipitation from lysates of *Dictyostelium* Nap1/Scar double KO cells expressing GFP-NAP1 and either Scar FL or ScarΔVCA, probed with anti-Nap1, anti-Pir121, anti-Scar and anti-Abi antibodies. **(D)** Nap1/Scar double KO cells expressing GFP-Nap1 (cyan) were transfected with LifeAct-mRFPmars2 (magenta) and Scar FL or ScarΔVCA as indicated, and imaged while migrating under agarose up a folate gradient. Representative cells are shown. The inset depicts a zoomed area used for the quantification of intensity of GFP-Nap1. Graphs on the side show the quantification of fluorescence intensity along the white line with 0 corresponding to the leading edge of the protrusion. The scale bar represents 10 µm. Error bars represent S.D. *n*> 17 cells counted for each condition. **(E)** Quantification of GFP-Nap1 recruitment as ratio between fluorescence intensity at the leading edge and in the cytosol. Bars show min to max values. **(F-H)** Scar KO cells were transfected with Scar FL or ScarΔVCA and allowed to migrate under agarose up a folate gradient while being observed by DIC microscopy at a frame interval of 3 seconds (1f/3s). Panels show quantification of speed **(F)**, Chemotaxis Efficiency Index (CEI, distance travelled in the direction of the gradient divided by the total distance travelled) **(G)** and directedness (distance between the beginning and the end divided by the total distance travelled) **(H)**. Data show means of > 30 analyzed fields from 7 independent experiments. Bars represent 5-95 percentile. For quantifications in **E, F, G** and **H** statistical significance was assessed by a two-tailed *t*-test. NS, not significant, ** p < 0.001, **** p < 0.00001.

### The Scar polyproline domain is essential for rescue of growth and motility in WASP-KO *Dictyostelium* cells

A direct actin polymerase activity of the WRC independent of the Scar/WAVE VCA domain has previously been linked to its polyproline domain^13,14^. Having found strong evidence that the WRC can promote actin polymerization independently of VCA-Arp2/3, we explored the possible involvement of the polyproline domain by further deleting this region from ScarΔVCA C-terminus and expressing the resulting ScarΔPVCA in *Dictyostelium* Scar KO cells (Fig.4A; Supplementary Fig.1E). Deletion of the Scar polyproline domain did not affect its inclusion in the WRC, as confirmed by immunoprecipitation of the other members of the complex (Supplementary Fig.1F), nor did it further increase the instability of the complex compared to ScarΔVCA expressing cells (Supplementary Fig.1A-E). Moreover, both truncated proteins rescued pseudopod formation and complex localization, with no differences in the recruitment levels compared to Scar FL containing WRC (Fig.4B-C). The rescue of pseudopod formation also translated into comparable rates of cell migration, with all motility parameters restored to the same levels as in cells expressing ScarΔVCA (Fig.4D; Supplementary Fig.1H-I). Thus, Scar is, even when lacking both the VCA and polyproline domains, able to promote pseudopod formation and restore cell motility of Scar KO cells.

In the protrusions of cells rescued with Scar FL, the Arp2/3 complex is clearly visible as a broad patch that extends behind the narrow leading edge localization of the WRC complex. Surprisingly, Arp2/3 is still recruited to the protrusions of cells expressing the truncated forms of Scar (Supplementary Fig.1G). To explain this difference between the B16-F1 and *Dictyostelium* cells, we investigated the possible involvement of other nucleating promoting factors (NPFs). Indeed, in *Dictyostelium* Scar KO cells WASP is able to replace Scar at the leading edge of protrusions and drive pseudopod formation and Arp2/3 complex activation^10,22,23^. Hence, we wondered if WASP could be responsible for the formation of pseudopods in cells expressing the truncated forms of Scar.

To avoid WASP complementation, we employed a Scar^DOX^/WASP KO system, an inducible double-knockout cell line whose Scar expression depends on the presence of doxycycline^10^. As previously described, when deprived of doxycycline these cells are unable to grow, to form pseudopods or migrate^10^. Scar^DOX^/WASP KO were transfected with Scar FL or the truncated forms of Scar, deprived of doxycycline for 48 hours to remove native Scar (Supplementary Fig.1J) and finally tested for rescue of growth and cell motility.

Expression of either Scar FL or ScarΔVCA rescued the formation of pseudopods, with the complex localized at the leading edge followed by enriched patches of F-actin. To the contrary, cells transfected with ScarΔPVCA were unable to form protrusions and could not move, similarly to untransfected cells deprived of doxycycline (Fig. 4E-F). Both Scar FL and ScarΔVCA fully rescued the growth rate of Scar^DOX^/WASP KO cells, while ScarΔPVCA expressing cells were unable to grow (Fig.4G). Moreover, ScarΔPVCA could not rescue Scar^DOX^/WASP KO migratory defect either. Conversely, ScarΔVCA expressing cells were able to migrate, with rescue of both speed, chemotaxis efficiency index and directionality (Fig.4H-I, Supplementary Fig.1K). Therefore, even without WASP compensation, the WRC complex lacking its VCA domain can promote pseudopod formation and rescue growth and motility. However, this ability is abolished after further deletion of Scar’s polyproline domain. In Scar KO cells, the deletion of Scar polyproline domain is compensated by WASP, which takes over Scar role and activates the Arp2/3 complex at the leading edge. In support to this hypothesis WASP was found to localize at the protruding leading edge of cells expressing either ScarΔVCA or ScarΔPVCA, while not in cells rescued with Scar FL (Fig.4J).

**Fig. 4.**
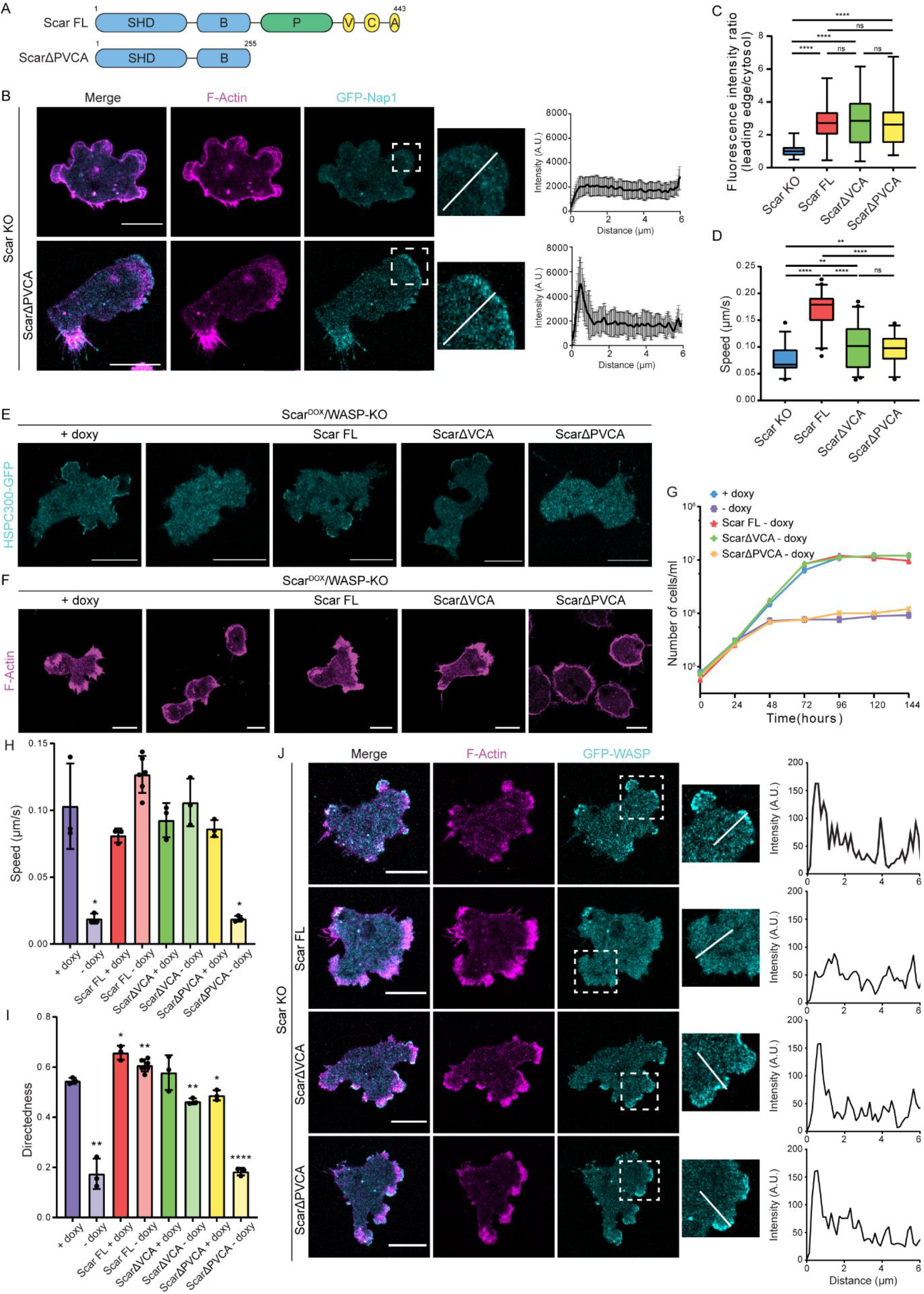
ScarΔPVCA is unable to rescue the formation of pseudopods in Scar/WASP KO *Dictyostelium* cells. **(A)** Schematic of Dictyostelium Scar FL and ScarΔPVCA showing amino acid numbers and domains. SHD – Scar homology domain; B – basic domain; P – polyproline domain; V – verprolin homology region; C – central region; A – acidic region. **(B)** Nap1/Scar double KO cells expressing GFP-Nap1 (cyan) were transfected with LifeAct-mRFPmars2 (magenta) and ScarΔPVCA, and imaged while migrating under agarose up a folate gradient. Representative cells are shown. The inset depicts a zoomed area used for the quantification of intensity of GFP-Nap1. Graphs on the side show the quantification of fluorescence intensity along the white line with 0 corresponding to the leading edge of the protrusion. The scale bar represents 10 µm. Error bars represent S.D. *n*> 17 cells counted for each condition. **(C)** Quantification of GFP-Nap1 recruitment as ratio between fluorescence intensity at the leading edge and in the cytosol. Bars show min to max values. **(D)** Scar KO cells were transfected with Scar FL, ScarΔVCA or ScarΔPVCA and allowed to migrate under agarose up a folate gradient while being imaged at a frame interval of 3 seconds (1f/3s). Panel shows quantification of speed. **(E-F)** Scar^DOX^/WASP KO cells were transfected with HSPC300-GFP (cyan in **E**) or LifeAct-mRFPmars2 (magenta in **F**) and rescued with Scar FL, ScarΔVCA or ScarΔPVCA as indicated. After growth in axenic medium supplemented or not with 10 µg/ml doxycycline for 48 hours, cells were imaged while migrating under agarose up a folate gradient for analysis of pseudopod formation. Representative cells are shown. The scale bar represents 10 µm. **(G)** Scar^DOX^/WASP KO cells were transfected with Scar FL, ScarΔVCA or ScarΔPVCA, grown in 6-well plates in axenic medium and counted every 24 h. Cells were starved of doxycycline at time = 0 h. Cells grown in the presence of doxycyxline were used as control. Data show a representative experiment from three independent experiments. **(H-I)** Scar^DOX^/WASP KO cells were transfected with Scar FL, ScarΔVCA or ScarΔPVCA. After 48 hours of doxycyxline starvation, cells were allowed to migrate under agarose up a folate gradient while being observed by DIC microscopy at a frame interval of 3 seconds (1f/3s). Panels show quantification of speed **(H)** and directednes **(I). (J)** Scar KO cells were transfected with LifeAct-mRFPmars2 (magenta) and GFP-WASP (cyan), and imaged while migrating under agarose up a folate gradient. Representative cells are shown. The inset depicts a zoomed area used for the quantification of GFP-Nap1 intensity. Graphs on the side show the quantification of fluorescence intensity along the white line with 0 corresponding to the leading edge of the protrusion. The scale bar represents 10 µm. For quantifications in **C, D, H** and **I** statistical significance was assessed by a two-tailed t-test. NS, not significant, * p, 0.05, ** p < 0.001, **** p < 0.00001.

### At least one polyproline domain is required to promote the formation of lamellipodia

We have shown that Scar/WAVE polyproline domain plays a direct role in actin polymerization in *Dictyostelium* cells. Similar activity was previously ascribed to the yeast WASP homolog Las17^14,15^, suggesting a possible conservation throughout evolution. We then moved back to B16-F1 cells. Following the same approach used for the VCA domain, we expressed WAVE2 without VCA and PP domains (referred to as WAVE2ΔPVCA, Supplementary Fig.2A) in WAVE1/2 KO cells. WAVE2ΔPVCA expression was confirmed by western blot, and it was comparable to that of WAVE2ΔVCA and WAVE2 FL (Fig.5A). Moreover, deletion of the polyproline domain did not affect the interaction with the other members of the complex, as verified by immunoprecipitation (Supplementary Fig.2B). We detected no differences in the migratory phenotype of WAVE2ΔVCA and WAVE2ΔPVCA. Despite being deprived of WAVE2 PP domain, cells expressing WAVE2ΔPVCA were still able to form lamellipodia at rates comparable to WAVE2ΔVCA (Fig. 5B-C). The deletion of WAVE2 polyproline domain also didn’t impact the recruitment of the complex to the lamellipodial protruding edge, where it accumulated in a narrow line along the plasma membrane similarly to WAVE2 FL and WAVE2ΔVCA-containing complexes (Fig.5E, Supplementary Fig.2C). The recovery of lamellipodia formation translated again in an increase of cell motility, with rescue of average speed to comparable levels to WAVE2 FL (Fig.5D). Based on these data, the WAVE2 polyproline domain did not appear to be essential for actin polymerization in B16-F1 cells.

**Fig. 5.**
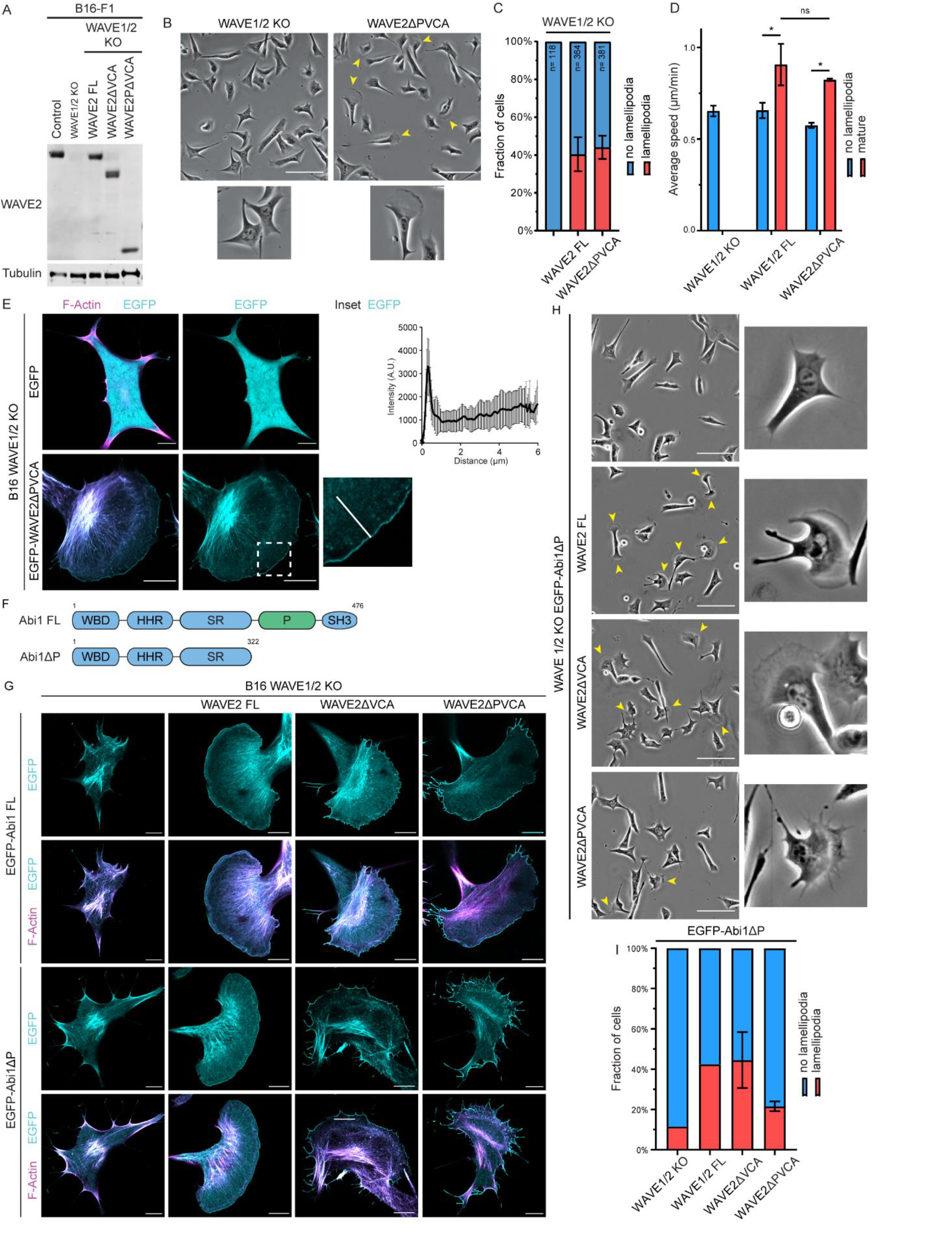
WAVEΔPVCA expression rescues lamellipodia formation in B16-F1 cells, but only when Abi’s polyproline domain is present. **(A)** Representative western blot of lysates of B16-F1 cells, WAVE1/2 KO cells, as well as KO cells expressing WAVE2 FL, WAVE2ΔVCA or WAVE2ΔPVCA to detect expression levels of WAVE complex components, as indicated. Tubulin was used as loading control. **(B)** Rescue of lamellipodia formation in WAVE1/2 KO cells transfected with WAVE2ΔPVCA and plated on laminin-coated 6-well plates. Yellow arrows indicate lamellipodia protrusions and insets below show magnifications of single cells. Scale bar = 100 µm. **(C)** Quantification of cells in **B** presenting with or without lamellipodia as a percentage. Error bars represent S.D., *n* > 118 cells counted for each condition from 2 independent experiments. **(D)** Random migration assay with WAVE1/2 KO cells expressing WAVE2 FL or WAVE2ΔPVCA, and analyzed as described in methods. Cells with and without lamellipodia are displayed separately. Graph shows mean values from 3 independent experiments. Error bars represent S.D. Statistical significance was assessed by a two-tailed t-test. NS, not significant, * p < 0.05. **(E)** WAVE1/2 KO cells were transfected with LifeAct-TagRed (magenta) and EGFP or EGFP-WAVE2ΔPVCA (cyan) as indicated, and plated on laminin-coated coverslips for analysis of lamellipodia morphology and localization of the WRC. Representative cells are shown. The inset depicts a zoomed in area used for the quantification of intensity of EGFP-WAVE2. The graph on the side shows the quantification of fluorescence intensity along the white line with 0 corresponding to the leading edge of the protrusion. The scale bar represents 10 µm. Error bars represent S.D., *n*> 21 cells counted for each condition from 3 independent experiments. **(F)** Schematic of mouse Abi1 FL and Abi1ΔP showing amino acid numbers and domains. WBD – WAVE binding domain; HHR – Homeodomain homology region; SR – serine rich region; P – polyproline domain; SH3 – SRC Homology 3 Domain. **(G)** WAVE1/2 KO cells were transfected with LifeAct-TagRed (magenta) and EGFP-Abi1 FL or EGFP-Abi1ΔP (cyan), and rescued with the WAVE2 constructs as indicated. Cells were plated on laminin-coated coverslips for analysis of lamellipodia morphology and localization of the WRC. Representative cells are shown. The scale bar represents 10 µm. **(H)** Rescue of lamellipodia formation in WAVE1/2 KO cells transfected with EGFP-Abi1ΔP and WAVE2 FL, WAVE2ΔVCA or WAVE2ΔPVCA as indicated. Cells were plated on laminin-coated 6-well plates and left to adhere for 6 hours before acquisition. Yellow arrows indicate lamellipodia protrusions and insets show magnifications of single cells. Scale bar = 100 µm. **(I)** Quantification of cells in **H** presenting with or without lamellipodia as a percentage. Error bars represent S.D., *n* > 100 cells counted for each condition.

We then decided to look for other polyproline domains that might still promote actin polymerization in cells expressing WAVE2ΔPVCA. The WRC complex contains two polyproline domains: one from Scar/WAVE and a second one found in Abi. A recent paper analyzed the effects of Abi1 knockout and demonstrated that the protein is able to recruit profilin-1 and N-WASP through its polyproline domain^24^. Thus, Abi polyproline domain could play a role similar to Scar/WAVE PP domain in promoting actin polymerization. To test this hypothesis we simultaneously removed the polyproline stretches in both Scar/WAVE and Abi by the generation of constructs for the expression of Abi full length (Abi FL) or Abi lacking its C-terminus (AbiΔP) (Fig.5F). In *Dictyostelium* cells, replacement of both native Scar and Abi with ScarΔPVCA and AbiΔP led to a highly unstable WRC, making impossible to draw any conclusion about their phenotype (Supplementary Fig.2D-E). In contrast, B16-F1 cells tolerate the deletion of both WAVE2 and Abi1 polyproline domains, allowing us to investigate the effects of their loss on cell migration and lamellipodia formation. Overexpression of EGFP-Abi1 and EGFP-Abi1ΔP was sufficient to outcompete the endogenous protein, so they were included in functional WRC localized at the leading edge of lamellipodia, as confirmed by immunoprecipitation (Supplementary Fig.2F) and live cell microscopy (Fig.5G). Deletion of Abi1 PP domain alone had no effects on cells ability to form lamellipodia (Fig.5G-H). Moreover, even when co-expressed with WAVE2ΔVCA, cells were still able to generate lamellipodia and rescued their migration speed (Fig.5H-I, Supplementary Fig.2G)). In contrast, expression of both WAVE2ΔPVCA and Abi1ΔP resulted in a nonfunctional WRC: the resulting cells could form only very sporadic protrusions which appeared to be extremely irregular and tipped by several filopodia (Fig.5G-I). Furthermore, this residual functionality could also be caused by the inclusion in the WRC of the endogenous full length Abi1. The observed defects in the generation of protrusions translated in the inability of these cells to efficiently migrate. Even the cells that were able to make protrusions had no significant increase in their speed when compared against the WAVE1/2 double knockout backgroud (Supplementary Fig.2G). Therefore, we conclude that at least one polyproline domain, from either Scar/WAVE or Abi, is required for the WRC to function in the absence of the VCA domain in B16-F1 melanoma cells.

## Discussion

Scar/WAVE’s function is fundamental in cell biology. Eukaryotic cells use broad protrusions called pseudopods or lamellipodia to migrate in 2D, which are based on polymerised branched actin networks. In normal cells, essentially all lamellipodia are initiated by signals activating the WRC. Cells without a functional complex, for example due to deletion of any subunit, do not make protrusions at all, their cytoskeleton and shape are profoundly aberrant, and their movement is compromised^9,10,25^. Three regions at the C-terminus of Scar/WAVE itself form the VCA domain, consisting of an actin-binding WH2 domain (the ‘V’ region), an acidic domain that recruits the Arp2/3 complex (the ‘A’ region), and a central helix (the ‘C’ region) that is believed to hold the entire WRC in an inactive state by a reversible intramolecular interaction^26^. Together the V, C and A regions can activate Arp2/3 and catalyze the production of new actin branched protrusions^27^.

For years it has been believed that the WRC works purely by activating the Arp2/3 complex^6,28^, with its VCA domain being considered the primary functional part, while other domains contribute only to its regulation^5,29,30^. However, in this study we demonstrate that Scar/WAVE lacking its VCA region still catalyzes the formation of actin protrusions in both B16-F1 melanoma and *Dictyostelium* cells and its expression is able to rescue the phenotype of WAVE1/2 or Scar KO cells, respectively. This result strongly contradicts the currently accepted dogma. The only precedent for this is in a recent study from the Weiner lab^16^. They demonstrated that the WRC can promote lamellipodia formation even in the absence of Arp2/3, while WAVE-null cells cannot. Similarly, here we show that WAVE2ΔVCA activity is uncoupled from Arp2/3: loss of the VCA domain stops WAVE2 from co-precipitating the Arp2/3 complex, showing that WRC containing WAVE2ΔVCA does not have an alternative pathway to bind it, and strongly reduces Arp2/3 recruitment to the leading edge. However, in these conditions F-actin still accumulates behind WAVE at the leading edge of protrusions. Hence, our data demonstrate that the WRC holds the ability to directly catalyze actin polymerization and suggest that it plays a more central role in forming lamellipodia than the Arp2/3 complex.

Until recently the field has focused only on the VCA domain of WASP-family proteins, almost completely ignoring the polyproline domain which was regarded as a passive linker^8^. However, many pieces of evidence accumulated over the last decade have begun to point out the importance of this region^13-15^. In the WRC both Scar/WAVE and Abi contain extensive proline-rich strings. Their sequences are not conserved in detail and vary in length, but their position within the protein and features, such as repeats of XPPPP, are consistent. In *Dictyostelium*, we have previously shown that the polyproline domain of Abi is dispensable for the activity of the WRC^31^. On the contrary, when the compensatory effect of WASP was abolished, the deletion of polyproline as well as the VCA domain of Scar made the WRC nonfunctional and unable to rescue pseudopod formation. Surprisingly, in B16-F1 cells WAVE2ΔPVCA expression was still able to rescue lamellipodia formation and motility. The reason for this difference could be the different stability of the truncated WRC in the two systems: in *Dictyostelium* deletion of the VCA domain strongly reduced the complex stability, while no similar effects were observed in B16-F1 cells. However, the situation changed when we simultaneously removed the polyproline stretches of both WAVE and Abi. In *Dictyostelium* the resulting WRC was even more unstable and the lack of stable complex made it impossible to investigate its function. In B16-F1 cells loss of both polyproline domains didn’t affect the complex stability, but it impaired the ability of the WRC to nucleate actin and rescue lamellipodia formation. The resulting cells could only form rare and aberrant protrusions which were unable to rescue cell motility. Taken together these results strongly suggest that polyproline domains are essential players in actin polymerization at protrusions, and at least one from either Scar/WAVE or Abi is required for the WRC to function.

This work poses new questions for the field. The molecular mechanism used by polyproline domains to promote actin polymerization is still unknown and requires further investigation. We know that these regions contain multiple weak G-actin binding sites which have been shown to be important for actin nucleation: by increasing the local concentration of actin, they can promote conditions for rapid polymerization by Arp2/3^32^. Moreover, in yeasts the binding of monomers by the PP domain of Las17 has been proved to allow actin nucleation in the absence of Arp2/3^15^. This property was also suggested serving to generate “mother” *de novo* filaments needed for the polymerization of branched actin networks by Arp2/3. However, this mechanism is poorly efficient and doesn’t explain how a WRC depleted of the VCA domain could completely rescue the formation of actin protrusions. Proline-rich regions are also binding sites for several proteins involved in actin nucleation^33^. Profilin can interact with G-actin as well as the PP domain of many proteins, including WASP-family members, and by doing so it recruits actin monomers to favor actin polymerization^34^. Other proteins interact with polyproline domains through EVH1, SH3 and WW domains^35-37^. Between them of particular interest for the generation of actin protrusions are VASP and BAIAP2. The first is a well-known actin polymerase involved in the assembly of actin filaments in sheet-like protrusions, filopodia, stress fibers and focal adhesions^38^. The second is a member of the I-BAR domain family of curvature-sensitive proteins that localizes with the WRC at lamellipodia and sites of saddle curvature^16^. Mass spectrometry analysis of WRC immunoprecipitation shows that both proteins are pulled down abundantly by WAVE2ΔVCA, but not by WAVE2ΔPVCA (data not shown). If any of them is the key transducer responsible for lamellipodia formation or if the WRC acts as a general hub that locally concentrates and promotes the interaction of a number of molecules needed for actin polymerization, like observed in clathrin structures^39^ and filopodia^40^, is still unknown. Lastly we cannot exclude the involvement of some other processes that do not depend on binding partners, such as direct membrane deformation at the leading edge and/or phase separation^41^.

Cells simultaneously assemble, maintain and disassemble actin filaments to continuously generate new actin protrusions. This complexity requires the ability to rapidly and efficiently turn on and off the actin polymerization machinery^4^. For this reason the manipulation of the lifetime of the active WRC is extremely important, as has been shown in simpler systems like WASP^42^. The inactive WRC is stable, but on activation it is subjected to exceptionally rapid proteolysis^43^. The WRC deleted of its VCA domain is deprived of its normal autoinhibition. Being in a “forced” open conformation, the complex is continuously subjected to proteolysis. Indeed, in *Dictyostelium* cells the deletion of Scar VCA domain alone is enough to strongly destabilize the WRC. On the contrary, B16-F1 cells have exceptionally low rates of degradation of the active WRC: even the deletion of both WAVE and Abi polyproline domains didn’t affect the complex stability. This property appears to be specific to this cell line and could explain why these cells are so strongly polarized. Identifying what makes these two systems so different could help deepen our knowledge of the mechanisms behind WRC removal and would allow a much cleaner understanding of pseudopods dynamics.

In conclusion, we propose the existence of a second pathway for actin polymerization mediated by the WRC which is independent of the Arp2/3 complex and requires the presence of at least one polyproline domain from either Scar/WAVE and Abi. Significant questions remain as to the different contribution of the two mechanism to the generation of cell protrusions. Moreover, our results are consistent in both B16-F1 mouse melanoma and *Dictyostelium* cells, ensuring our conclusions are general. Thus, polyproline regions may play a similar role in the other WASP family proteins.

## Online Methods

### Antibodies and constructs

Antibodies and DNA constructs are listed in Supplementary Tables 1 and 2, respectively.

### DNA constructs

All primers are listed in Supplementary Table2.

WAVE2ΔVCA and WAVE2ΔPVCA were amplified by PCR from pSP330 and cloned into pCDNA3.1 or pEGFPC1 vector using KpnI/XbaI. The coding sequence of mouse Abi1 was amplified from cDNA (ID:3498068, Dharmacon) and cloned into pEGFPC1 using BglII/SalI. *Dictyostelium* Scar was amplified by PCR from genomic DNA and cloned in pDM304 expression vector^44^. Scar co-expression constructs were generated by ligating pSP149, pSP259, pDM459 or pAD58 NgoMIV fragments into Scar expression vectors.

The fidelity of all constructs was verified by sequencing.

### Mammalian cell lines and growth conditions

Cell lines are listed in Supplementary Table 3.

Mouse melanoma B16-F1 cells and derived WAVE1/2 double knockout clones were cultured in Dulbecco’s Modified Eagles Medium (DMEM, Gibco) supplemented with 10% FBS (Gibco) and 2mM L-glutamine (Gibco), and maintained in 10cm plastic Petri dishes at 37ºC and under 5% CO_2_.

### Transfection of mammalian cell lines

For WB and imaging experiments, B16-F1 cells were plated on a 6-well plate, grown to 70% confluency and later transfected with Lipofectamine 2000 following the manufacturer’s guidelines with 2-5 µg DNA. For GFP-trap experiments, cells were plated on 15cm plastic Petri dishes, grown to 70% confluency and later transfected with Lipofectamine 2000 following the manufacturer’s guidelines with 10 µg DNA.

### Immunofluorescence staining and analysis

Cells were seeded onto sterile 13mm glass coverslips or on 96-well glass bottom dishes with black well walls (CellCarrier Ultra, PerkinElmer) that had been previously coated with 10µg/ml laminin (Merk) diluted in sterile PBS and allowed to adhere and form lamellipodia overnight. The next day cells were fixed with 4% paraformaldehyde for 20 min, RT. Samples were then permeabilized with 0.2% Triton X-100 for 10 min and washed three times for 5 min in PBS/0.1M glycine before incubation with blocking buffer (5% BSA, PBS) twice for 15 min. Primary antibodies were diluted in blocking buffer and incubated with coverslips in a dark and humidified chamber overnight. Samples were washed six times in PBS + 0.5% BSA and incubated with the secondary antibodies for 45 min in a dark, humidified chamber at room temperature. Samples were then incubated with phalloidin, CellMask and/or DAPI for 30 min at RT. Coverslips were washed twice in PBS + 0.5% BSA, three times in PBS and once in MilliQ water before mounting with ProLong Glass Antifade Mountant with NucBlue (Invitrogen). Samples for HSC were washed and kept in PBS.

Imaging was conducted on a Zeiss LSM 880 Airyscan confocal microscope or on an Opera Phenix HighContent Screening System (PerkinElmer).

For fluorescence intensity measurements of ArpC2 and phalloidin stainings, lamellipodia regions were encircled using Fiji software (ImageJ), and an extracellular region defined as background. Average intensities of background regions were then subtracted from average fluorescence intensities derived from lamellipodia regions.

### High Content Screening

Cells were imaged using an Opera Phenix high-throughput microscope and multiparametric image analysis carried out using Harmony 4.9 software’s supervised machine learning tool, Phenologic (both Perkin Elmer). Thirty fields of view per well were acquired, at 20x magnification, with 4 planes covering the depth of the cell and analysed in maximum projection. GFP positive cells were selected, and a small training population of these manually subdivided into 3 morphological classes. Using multiple measurements of morphology and actin distribution, the software then segregated the whole population on this basis. Each morphology was then expressed as a percent of total cells.

### Live cell imaging

B16-F1 cells were seeded on glass-bottom laminin-coated dishes for at least 4-5 hours and then imaged using a Zeiss LSM 880 Airyscan confocal microscope with a heated incubator with a 63x/1.40 NA objective. Images were acquired using the ZEN imaging software every 30 sec for 10 min at 37ºC and under 5% CO_2_.

### Random migration assay for mammalian cells

Six-well glass-bottom plates were coated overnight as described above. Cells (1 × 10^5^) were plated and, after 4-5 hours, imaged every 10 min for 17h using a Nikon TE2000 microscope with a Plan Fluor 10x/0.30 objective and equipped with a CO_2_ perfused chamber heated at 37ºC. For analysis, individual cells were manually tracked using Fiji software (ImageJ).

### *Dictyostelium discoideum* cells

Cell lines are listed in Supplementary Table 3.

The axenic *Dictyostelium discoideum* strain Ax3 was used as the WT. All strains were grown in HL5 medium (ForMedium) with 100µg/ml penicillin-streptomycin (Gibco) in 10cm plastic Petri dishes and incubated at 21ºC.

### Transfection of *D. discoideum*

Some 1.0×10^7^ cells per transfection were first centrifuged (3min, 340*g*, 4 ºC), washed with 10 ml ice-cold electroporation buffer (E-buffer: 5mM Na_2_HPO_4_, 5mM KH_2_PO_4_ and 50mM sucrose) and resuspended in 420µl ice cold E-buffer. Cells were transferred into ice-cold 0.2cm electroporation cuvette and electroporated with 5-7µg extrachromosomal plasmid at 500V using ECM399 electroporator (BTX Harvard apparatus), giving a time constant of 3-4ms. Transfected cells were transferred into HL5 medium, including glucose, vitamins and microelements (ForMedium) in 10cm plastic Petri dishes. After 24 hours, transfectants were selected and maintained using 50µg/ml hygromycin or 20µg/ml G418.

### *D. discoideum* under-agarose chemotaxis assay

Cell migration and lamellipodia formation were examined by under agarose folate chemotaxis assay^45,46^. The surface of 6-well glass-bottom dishes (MatTek) were coated with 5% BSA for 10 min, washed with distilled water and let air-dry. Then, 0.4% w/v SeaKem GTG Agarose was melted in LoFlo medium (Formedium). After cooling of agarose, 10µM folate was added and the mix was poured into coated dishes and allowed to set. A 5mm wide well was cut in the agarose using a scalpel and 2 × 10^6^/ml cells were seeded inside. After 3-4 hours, cells were imaged by phase contrast microscopy with a Nikon Eclipse TE2000-E inverted microscope system equipped with a QImaging RETIGA EXi FAST 1394 CCD camera and a pE-100 LED illumination system (CoolLED) at 525nm. A 10x/0.45 NA Ph1 objective was used. Images were taken at 100 sec intervals for 2 hours and imaging was controlled using Micro-Manager software. Chemotactic parameters were calculated using home-made plug-in of ImageJ written by Dr. Luke Tweedy^9^.

For cells expressing fluorochrome-tagged proteins, 0.5% w/v SeaKem GTG Agarose was melted in LoFlo medium (Formedium) and was poured into BSA-coated 50mm glass-bottom dishes (MatTek). Once set, the agarose was cut using a scalpel to create two wells separated by a 5mm bridge. About 2 × 10^6^/ml cells were seeded on the left well, while the other well was filled with 100µM folic acid diluted in LoFlo. After 3-4 hours cells were imaged using a Zeiss LSM 880 Airyscan confocal microscope with a 63x/1.40 NA objective. Images were acquired using the ZEN imaging software every 3 sec.

All microscopy was carried out at room temperature.

### *Dictyostelium* cell growth assay

Cells were plated on 60-well dishes and counts were performed every 24 hours with a CASY Cell Counter (OMNI Life Science).

### GFP-trap pull down

For B16-F1, cells growing on a 15 cm dish were washed in ice-cold PBS twice. Lysis was performed by scraping cells in Lysis Buffer (25mM Tris-HCl pH7.5, 100mM NaCl, 5mM MgCl_2_, 0.5% NP40, 1X Halt protease and phosphatase inhibitor cocktails). Lysates were collect in ice-cold tubes and kept in ice for 1 hour. Tubes were then centrifuged for 15 min at 13000 rpm 4ºC. The supernatants were transferred to a clean Eppendorf tube and measured for protein concentration using Precision Red (Cytoskeleton). A volume of 25 µl of GFP-trap beads (Chromotek) were equilibrated following manufacturer’s protocol. Equal amounts of lysates were mixed with the beads and incubated on rotation for 2 hours at 4ºC. Beads were spun down (2500*g*, 4ºC, 2 min) and washed three times in Wash Buffer (25mM Tris-HCl pH 7.5, 100 mM NaCl, 5mM MgCl_2_, 1X Halt protease and phosphatase inhibitor cocktails). Samples were eluted after incubation with 2X NuPAGE LDS sample buffer and heated for 10 min at 70 ºC before loading for SDS-Page.

For *Dictyostelium*, cells growing on a 15 cm dish were collected in 0.017 M Soerensen Na-K phosphate buffer pH 6.0 and spin down. Pellets were lysed by resuspending them in *Dictyostelium* Lysis Buffer (50mM Tris-HCl pH 8.0, 100mM NaCl, 30mM MgCl_2_, 0.1% Triton X-100, 1X Halt protease and phosphatase inhibitor cocktails, 1mM DTT) and kept in ice for 30 min. Lysates were cleared by centrifugation (13000 rpm, 15 min, 4ºC). The supernatants were transferred to a clean Eppendorf tube and measured for protein concentration using Precision Red (Cytoskeleton). Lysates were mixed with pre-equilibrated GFP-trap beads as described above. Beads were washed three times in *Dictyostelium* Wash Buffer (Tris HCl pH 7.5, 150mM NaCl, 0.5mM EDTA, 1X Halt protease and phosphatase inhibitor cocktails). To elute the proteins from the beads, 2X NuPAGE LDS sample buffer was added and boiled (70 ºC 10 min). Protein samples were analysed by SDS-Page.

### SDS-Page and Western Blotting

For preparation of B16-F1 whole lysates, cells were washed with PBS and lysed by scraping cells in RIPA buffer (150mM NaCl, 10mM Tris-HCl pH 7.5, 1mM EDTA, 1% Triton X-100, 0.1% SDS, 1X Halt protease and phosphatase inhibitor cocktails), collecting in ice-cold tubes and keeping them in ice for 1 hour. Then tubes were centrifuged for 15 min at 13000 rpm at 4ºC. The supernatants were transferred to a clean Eppendorf tube and measured for protein concentration using Precision Red (Cytoskeleton).

*Dictyostelium* cells were lysed by directly adding 1X NuPAGE LDS Sample buffer (Invitrogen) containing 20mM DTT, followed by incubation at 70 ºC for 10 min.

The 40µg of protein lysate was resolved on NuPAGE Novex 4-12% or 12% Bis-Tris gels (ThermoFisher) and transferred onto a nitrocellulose membrane (Amersham Protran, Merk) at 100V for 1 hour. Membranes were blocked with 5% semi-skimmed milk diluted in TBST (10mM Tris pH 8.0, 150mM NaCl, 0.5% Tween-20) for 1 hour prior to overnight incubation with the primary antibody at 4ºC on a roller shaker. Membranes were then washed three times for 10 min each in TBST and incubated with secondary AlexaFluor conjugated antibodies for 1 hour at room temperature. The blots were washed again for 10 min in TBST three times before being imaged on the Li-Cor Odyssey CLx machine. Images were then analysed using Image Studio Lite Version 5.2 (LI-COR). For *Dictyostelium* samples MCCC1^47^ and for mammalian samples α-tubulin were used as a loading control.

### Quantification and statistical analysis

Data analyses were done using GraphPad Prism 8. Statistical tests used are indicated in each figure legend. p values < 0.05 were considered as significant: *: p < 0.05, **: p < 0.01, ***: p < 0.001, ****: p < 0.0001.

## Acknowledgements

We thank the Beatson Advanced Imaging Resource (BAIR) (A17196), in particular Margaret O’Prey, David Strachan and Nikki Paul for their training and assistance on the microscopes. We also acknowledge the CRUK Beatson Institute Core Services (Grant C596).

## Authors’ contribution

S.B. and R.H.I. conceived the study and designed the experiments. S.B. performed most of the experiments, analyzed the data and wrote the manuscript. S.P. generated plasmids and performed experiments. S.C., P.P., J.W. and P.T. assisted S.B. in experiments, analyzed data, and provided essential reagents. L.T. performed data and image analysis. L.M. performed HighContent Screening analysis. All authors have read and agreed to the published version of the manuscript.

## Data availability statement

Further information and requests for resources and reagents may be directed to Robert H. Insall and Simona Buracco.

