## Supplementary legends for "The Scar/WAVE complex drives normal actin protrusions without the Arp2/3 complex, but proline-rich domains are required"

- Supplementary Fig.1

**(A-E)** Relative levels of WRC subunits in WT AX3 cells and Scar KO cells transfected with Scar FL, ScarΔVCA and ScarΔPVCA. Cells were harvested during the growth phase and boiled in SDS sample buffer. Equal amounts of proteins were separated by SDS-PAGE and stained for WRC subunits using specific antibodies. **(A-D)** Graphs show normalized values of protein levels, as quantified by densitometry of 14 immunoblots for each subunit and cell line. Error bars indicate means ± S.D. Statistical significance was assessed by a two-tailed t-test against WT control. * $p < 0.05$; ** $p < 0.001$; *** $p < 0.0001$; **** $p < 0.00001$. **(E)** Representative western blot. MCCC1 was used as loading control. **(F)** Representative western blot of GFP immunoprecipitation from lysates of *Dictyostelium* Nap1/Scar double KO cells expressing GFP-NAP1 and either Scar FL, ScarΔVCA or ScarΔPVCA, probed with anti-Nap1, anti-Pir121, anti-Scar and anti-Abi antibodies. **(G)** Nap1/Scar double KO cells expressing GFP-Nap1 (cyan) were transfected with mRFPmars2-ArpC4 (magenta) and Scar FL, ScarΔVCA or ScarΔPVCA as indicated, and imaged while migrating under agarose up a folate gradient. Representative cells are shown. The scale bar represents 10 µm. **(H-I)** Scar KO cells were transfected with Scar FL, ScarΔVCA or ScarΔPVCA and allowed to migrate under agarose up a folate gradient while being observed by DIC microscopy at a frame interval of 3 seconds (1f/3s). Panels show quantification of CEI **(H)** and directedness **(I)**. Statistical significance was assessed by a two-tailed t-test. NS, not significant, * $p < 0.05$; ** $p < 0.001$; *** $p < 0.0001$; **** $p < 0.00001$. **(J)** Loss of both SCAR and WASP from $Scar^{DOX}$/WASP KO cells verified by Western blotting with anti-SCAR and anti-WASP antibodies. Substantial repression of SCAR is seen after 48 h. Cells are completely deleted for WASP; only SCAR is inducible. MCCC1 is the loading control.

- Supplementary Fig.2

**(A)** Schematic of mouse WAVE2 FL and WAVE2ΔPVCA showing amino acid numbers and domains. WHD – WASP homology domain; B – basic domain; P – polyproline domain; V – verprolin homology region; C – central region; A – acidic region. **(B)** Representative western blot of GFP immunoprecipitation from lysates of WAVE1/2 KO cells expressing either EGFP alone, EGFP-WAVE2 FL, EGFP- WAVE2ΔVCA or EGFP-WAVE2ΔPVCA, probed with anti-NCKAP1 and anti-Cyfip1. **(C)** WAVE1/2 KO cells were transfected with EGFP-tagged WAVE2 (cyan) constructs as indicated, and plated on laminin-coated coverslips for analysis of lamellipodia morphology and localization of the WRC. Graph shows quantification of

WAVE2 recruitment as ratio between fluorescence intensity at the leading edge and in the cytosol. Bars show min to max values. Statistical significance was assessed by a two-tailed *t*-test. NS, not significant. **(D)** Representative western blot of cell lysates of *Dictyostelium* AX3 WT, Scar and Abi single and double KO, as well as KO cells expressing Scar FL, ScarΔPVCA, Abi FL and/or AbiΔP as indicated to detect expression levels of WRC components. MCCC1 was used as loading control. **(E)** Representative western blot of GFP immunoprecipitation from lysates of *Dictyostelium* Scar/Abi double KO cells expressing GFP-Pir121 and either Scar FL, ScarΔPVCA, Abi FL and/or AbiΔP, probed with anti-Nap1, anti-Pir121, anti-Scar and anti-Abi antibodies. **(F)** Representative western blot of GFP immunoprecipitation from lysates of WAVE1/2 KO cells rescue with the indicated WAVE2 constructs and expressing either EGFP alone, EGFP-Abi1 or EGFP-Abi1ΔP, and probed with anti-NCKAP1 and anti-Cyfip1 to prove inclusion in the WRC. **(E)** Random migration assay with WAVE1/2 KO cells expressing EGFP-Abi1ΔP rescued with the indicated WAVE2 constructs, and analyzed as described in Methods. Cells with and without lamellipodia are displayed separately. Error bars represent S.D.

- Supplementary Table 1

List of antibody information regarding provider, catalogue number and working dilution of antibodies and fluorescent probes/stains used in this study.

- Supplementary Table 2

List of construct information regarding the plasmids and oligonucleotides used in this study.

- Supplementary Table 3

List of cell lines used in this study.
