## Supplementary figures and images for "The Scar/WAVE complex drives normal actin protrusions without the Arp2/3 complex, but proline-rich domains are required"

### Supplementary Fig.1

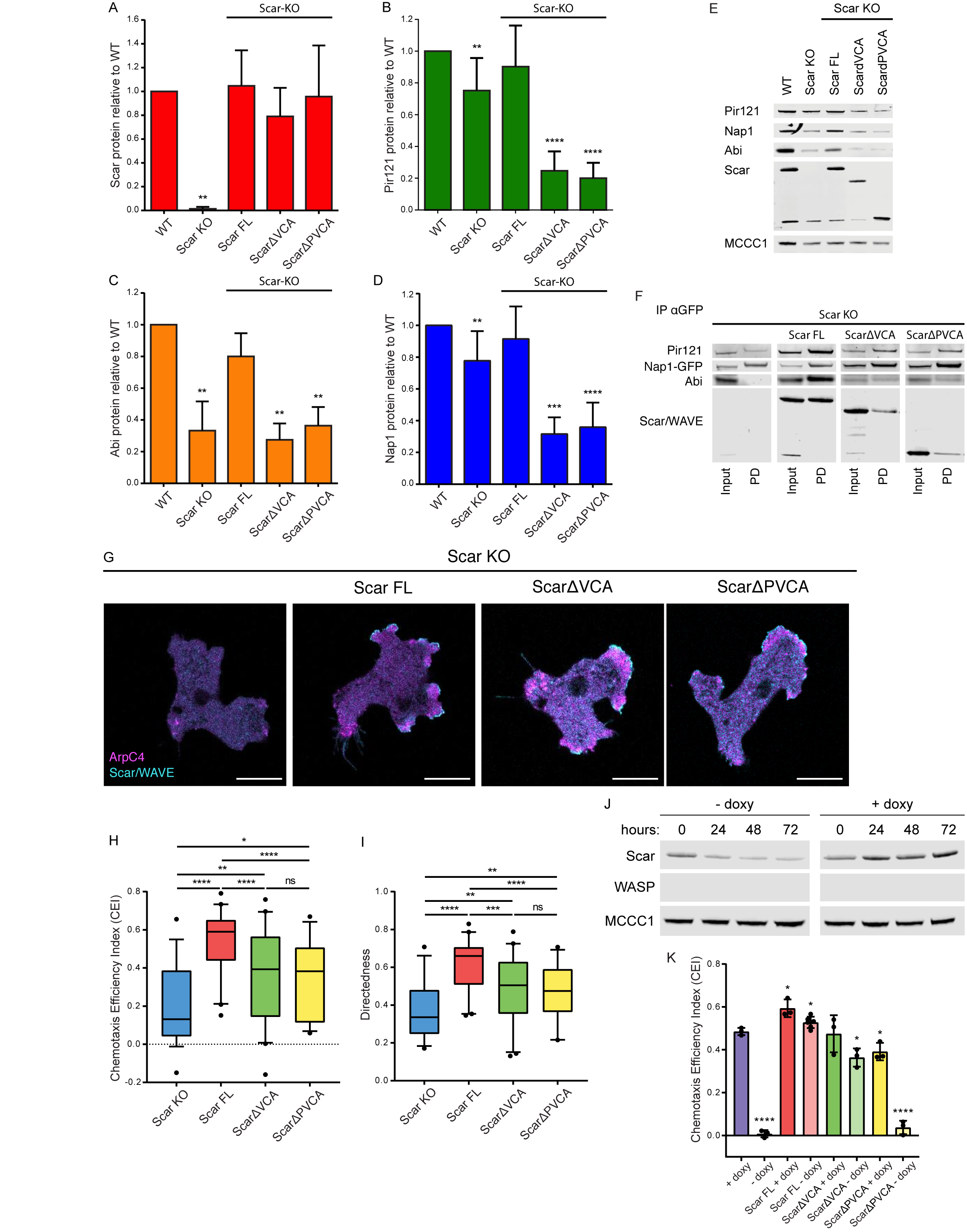

### Supplementary Fig.2

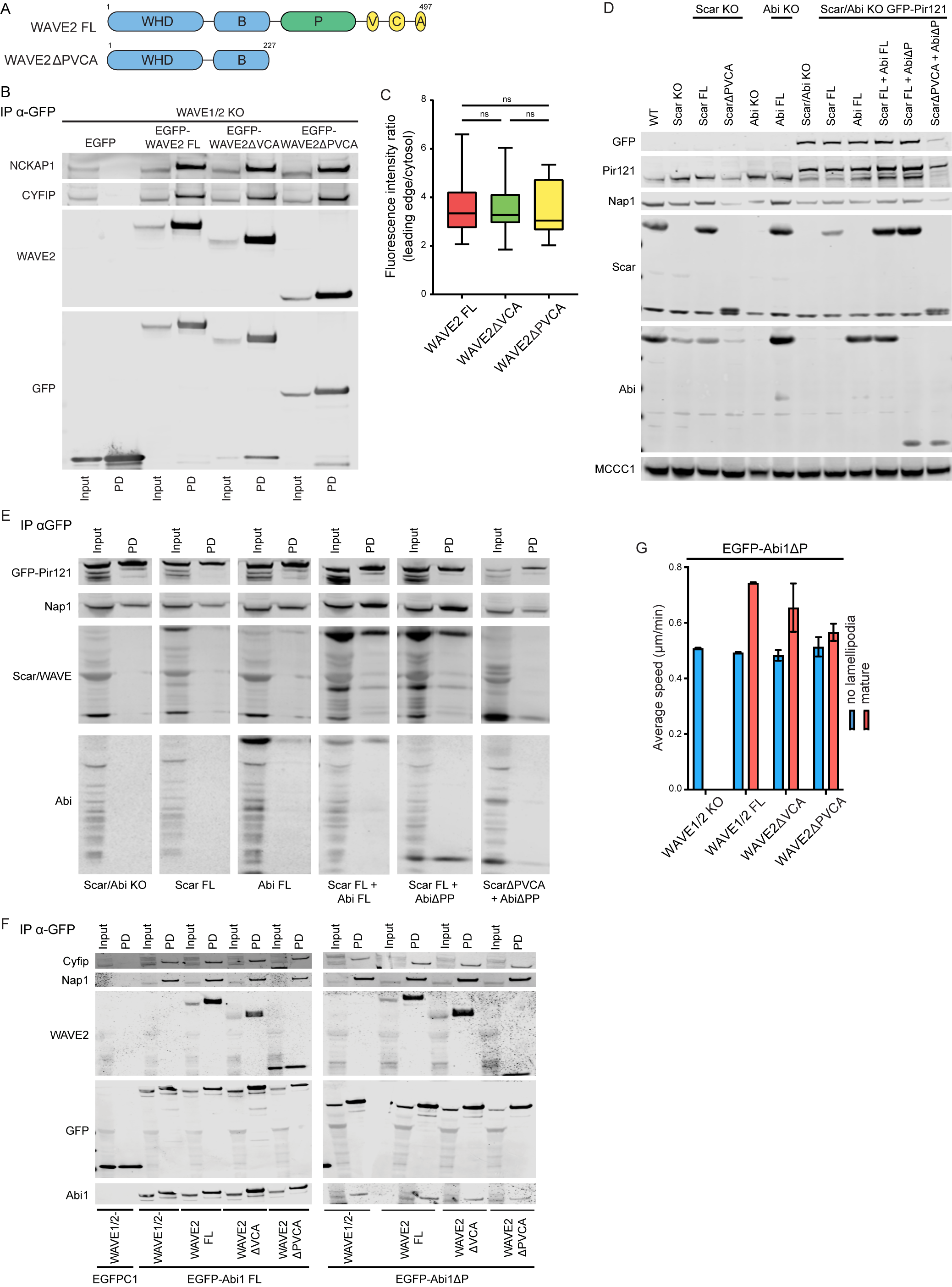
